# Using metabolic data to uncover the ontogeny of endothermy in an altricial bird

**DOI:** 10.1101/2023.08.29.555287

**Authors:** Elana Rae Engert, Fredrik Andreasson, Andreas Nord, Jan-Åke Nilsson

**Affiliations:** Department of Biology, Section for Evolutionary Ecology, Lund University, Ecology building, Lund S-22362, Sweden

**Keywords:** Metabolic rate, Development, Endothermy, Altricial birds, Cooling rate, Ecophysiology

## Abstract

Altricial songbirds transform themselves from naked poikilotherms to fully-feathered endothermic homeotherms over a matter of weeks from hatching to fledging. The timing of development of endothermy has been studied in altricial birds for over half a century. However, the potential determinants, constraints, and flexibility of the onset of endothermy are not yet fully understood. We experimentally investigated whether brood size influences the ontogeny of endothermic heat production in 4-8 day-old nestling blue tits (*Cyanistes caeruleus*) in southern Sweden. We found that both nestlings’ capacity to produce heat and the proportion of nestlings exhibiting a metabolic response to a cooling challenge (15 °C for 15 min) increased with age. However, we did not find an effect of brood size on the ontogeny of endothermy and this may be explained by the lack of differences in body mass of nestlings in enlarged and reduced broods in our study. The metabolic response in all age groups was brief and not sufficient to sustain a stable body temperature. Our study shows that the development and use of endothermy is present at an early age in nestling blue tits, but further study is needed to disentangle the factors contributing to individual variation in the timing of these developments. This will serve to improve our understanding of a developmental milestone of cold tolerance in birds in the temperate zone that not only face warmer average temperatures, but also increasingly frequent cold snaps and extreme weather.

**Summary Statement:** This study serves to improve our understanding of the ontogeny of endothermy in altricial birds while examining the potential role of brood size in the timing thereof.

## Introduction

Songbirds have altricial young that are small and featherless with little capacity to maintain a stable body temperature in the peri- and neonatal stages (Hohtola & Visser, 1998). Thus, they are poikilothermic, meaning that their body temperature closely follows ambient temperature (Pereyra & Morton, 2001). Nestlings of many bird species undergo a rapid development and become endothermic part-way through their nestling stage when they develop enough thermogenic tissue in combination with sufficient body size and insulative feathers (Andreasson et al., 2016; Olson, 1992; Węgrzyn, 2013).

Shivering in skeletal muscle is the main avenue for thermogenesis in birds (Dawson & Whittow, 2000; Hohtola, 2002). The thermogenic capacity of individual nestlings increases rapidly when they become capable of shivering thermogenesis (Morton & Carey, 1971), which is usually when they reach 50 – 70% of their adult mass (for a review see Dunn (1975)). However, if the development and use of thermogenic tissues consumes energy that could instead be directed to overall growth in body size, this would potentially result in an energy allocation tradeoff (Andreasson et al., 2016; Dawson & Evans, 1957, 1960). On the other hand, the transition of nestlings from poikilothermic to endothermic is closely linked to the time and effort parents allocate to different aspects of parental care. For example, endothermic nestlings no longer need to be brooded by parents, which frees up more time for parents to allocate more energy to other activities, such as foraging. This becomes more important as nestlings grow larger and have increased feeding requirements.

Due to the decrease in surface area to volume ratio of the brood with an increase in the number of nestlings, larger broods can achieve “effective homeothermy” as a unit faster compared to smaller broods (Dunn, 1975; Mertens, 1969; O’Connor, 1975; Sullivan & Weathers, 1992) and can have a more stable nest temperature during the nestling stage (Andreasson et al., 2016). However, nestlings in larger broods may face increased sibling competition for feedings (Nilsson, 2002; Stjernman et al., 2004) which can negatively affect body mass (Nilsson & Gårdmark, 2001). Disentangling the effects of brood size on the development of endothermy from the insulative properties of the brood requires studying the development of homeothermy in single nestlings.

Previous studies have focused primarily on body temperature cooling rate as a proxy for endothermy. However, cooling rates are influenced by different factors related to both growth (overall body size) and development (i.e., feathers and capacity for shivering) of nestlings. Thus, with temperature data alone, the different proximate explanations for endothermy cannot be separated. Alternatively, using metabolic data can reveal whether animals are actively producing heat to stay warm when ambient temperature drops, which is necessary to draw conclusions about the onset of endothermy (Hill, 2022). This represents a current knowledge gap that would serve to complement and clarify the conclusions we draw from cooling rate data alone.

In a previous study, we have focused on body temperature data to investigate the effect of brood size on the timing of homeothermy in individual nestlings (Andreasson et al., 2016). However, in the absence of metabolic data, we were missing a definitive link to endothermy. Now that it has become possible for us to bring respirometry measurement devices into the field, we can build on our previous work by incorporating metabolic data. To do this, we investigated the effect of brood size manipulations on the onset of endothermy in a small passerine, the blue tit (*Cyanistes caeruleus*), in southern Sweden. Blue tits are hole-nesting birds that have a clutch size of 10-14 eggs and a nestling period of about 3 weeks. Their relatively long nestling period and large clutch size (compared to open-cup nesting passerines; von Haartman, 1957) and their propensity to use nest boxes makes them a practical species for studying development in the nest and for brood size manipulations. We predicted that if the timing of developing endothermy is influenced by brood size, nestlings in experimentally enlarged broods would show a later onset of endothermy compared to reduced broods due to having less energy to put towards developing or using thermogenesis, and/or due to a reduced need for thermogenesis.

## Methods

The experiment was conducted in the Revinge area, surrounding Lake Krankesjön in southernmost Sweden (55°42′N, 13°28′E), between April and June 2022. The landscape is comprised of grasslands that are grazed by cattle, with patches of deciduous and coniferous forest. In this area, around 500 nest boxes have been monitored each breeding season since 1983. The nest boxes are constructed from wood, have an entrance hole (26 mm diameter) that is large enough for blue tits (*Cyanistes caeruleus*) and marsh tits (*Poecile palustris*), but too small for sympatric great tits (*Parus major*). Starting April 18^th^ 2022, nest boxes were checked weekly to monitor nest building and egg laying. The date of the first egg was back-calculated assuming that one egg was laid per day. Nests were checked daily for hatching starting 12 days after the last egg in the clutch was laid. The hatching date was considered as nestling day 0.

On day 3, when nestlings were old enough to be safely transported short distances, brood sizes were manipulated by moving four nestlings from one nest to another. Pairs of nests were chosen so that they were within a 15-minute drive of each other, had hatched on the same date and had between 9 and 14 nestlings. At each nest, nestlings were counted and all of them were weighed together in a cloth bag using a Pesola spring scale (Pesola AG, Baar, Switzerland) to the nearest 0.1 g. The four nestlings that were to be transferred were then removed from the reduced brood and weighed, transported, and added to the nest cup of the nest to be enlarged. Hatching dates did not differ significantly between enlarged and reduced broods (henceforth, E and R) and the mean hatch date overall was May 19 (t-test, t_24_ = -0.75, p = 0.46; E: mean ± SD = 48.4 ± 3.3 April day, R: mean ± SD = 49.4 ± 3.6 April day; April day is number of days since the 31^st^ of March). Brood size before manipulation did not differ between brood size categories (t-test, t_24_ = -0.05, p = 0.96; E: mean ± SD = 10.9 ± 0.4 nestlings, R: mean ± SD = 10.9 ± 0.3 nestlings). The mean nestling mass did not differ significantly between E (mean ± SD = 2.8 ± 0.4g) and R (mean ± SD = 2.7 ± 0.5g) broods pre-manipulation (t-test, t_24_ = 0.52, p = 0.61) and there was no difference in mass between the four transferred nestlings (mean ± SD = 2.8 ± 0.5g) and those remaining in the donor nest (mean = 2.7 ± 0.5g) in R broods (t-test, t_26_ = 0.49, p = 0.63). Nestlings that were moved between nests were marked with red nail polish on the toenails and ringed at age 7-9 days with an aluminum ring. All other nestlings were ringed on day 14, at which point we also measured the tarsus length to the nearest 0.01 mm, wing length to the nearest 0.5 mm, and body mass to the nearest 0.1 g for all nestlings. We caught and ringed parents on day 14.

Respirometry measurements were conducted when nestlings were 4, 6, and 8 days old. Only unmoved nestlings (i.e., those not marked with nail polish) were used for measurements. In all experiments, four nestlings were measured at the same time to get a representative sample of the brood in an efficient way. On each measurement day, we selected four nestlings from each nest, avoiding the smallest or largest ones in the brood. The nestlings were moved to the experimental setup, which was portable and placed within 50 – 100 meters of the nest to minimize disturbance to the feeding parents. The four nestlings were weighed together in a bag and then each nestling was placed into a wire basket separated into four sections to prevent nestlings from huddling. The wire basket was made from stainless steel (1.3x1.3 cm mesh) and spray-painted matte black (Figure S1). During all measurements, the chicks were contained in respirometry chambers made of glass with plastic locking lids and a rubber seal. The insides of the lids were covered with aluminum tape to minimize water vapor permeability. The inner surfaces of the lid and glass container were covered with matte black spray paint. Two sizes of respirometry chamber were used to accommodate nestlings of different ages; 600ml, 15×15×6cm (day 4 and 6) and 1000ml, 21×15×6cm (day 8).

We used an FMS (Field Metabolic System, Sable Systems, Las Vegas, NV, USA) to measure CO_2_ production and O_2_ consumption at 1 Hz. The FMS was calibrated directly before the start of the experiment and mid-way through (9 days later) the study. The CO_2_ analyzer was zero-calibrated using 100% N_2_ gas and span-calibrated using 0.5% CO_2_ in N_2_ gas. O_2_ was spanned to 20.95% in dry air (dried using Drierite®; Hammond Drierite Company, Xenia, OH, USA). Water vapor pressure (WVP) was measured simultaneously with CO_2_ and O_2_, but the lag time of the measurement precluded us from using the instantaneous WVP dilution correction for calculations of VO_2_ and VCO_2_. Consequently, we used wet measurements of CO_2_ and O_2_ for all calculations.

The respirometry chambers were placed in two portable electric cooling and heating boxes (24 L, 49-513, Biltema, Gothenburg, Sweden). We pulled dry air over the respirometry chambers at 350.0 ± 0.1ml / min (mean ± SD; standard temperature and pressure, STP) on day 4, 538.5 ± 21.1 ml / min on day 6, and 793.6 ± 10.0 ml / min on day 8. One of the boxes was set to 36°C (henceforth ‘warm’) to simulate a normal nest temperature (Andreasson et al., 2016; Visser, 1998) and the other was set to 15°C (henceforth ‘cold’) which is below the thermoneutral zone, in order to induce a cold response (Mertens, 1969) and for comparability to previous studies (Andreasson et al., 2016; Page et al., 2022). The temperature of each box was controlled using digital thermostats (KT3100, Ketotek, Xiamen, China). Temperature inside the respirometry chambers was measured continuously during measurements and was 35.8 ± 0.7°C in the warm chamber and 15.1 ± 0.8°C in the cold chamber (mean ± SD; measured using a type T (copper-constantan) 36G thermocouple). The thermosensitive junction of a similar thermocouple was taped in contact with the skin in the armpit of one nestling to continuously monitor skin temperature during the experiment.

Nestlings were given time to acclimate in the warm box for a mean ± SD of 8.0 ± 5.1 min before the first respirometry measurements were taken. Nestlings were first measured in the warm box for 9.6 ± 2.2 min, (mean ± SD), which was long enough for O2 levels to stabilize. Next, nestlings were measured for 15 minutes in the cold box, to measure the metabolic response to a cold challenge. In 6 pilot trials, nestlings were measured for 18-22 minutes to see if their O_2_ consumption would stabilize in the cold. Since a stable plateau in O_2_ consumption was not consistently reached within this time frame, we standardized recording time to 15 minutes, which was sufficient to capture the metabolic response to cooling. The mean ± SD time of transfer from the warm box to the cold box was 30.5 ± 12.5 sec (range = 19 – 84 sec). Baseline air measurements were taken for 6.5 ± 4.0 min before the warm session, and before and after the cold session (Figure 1).

**Figure 1.**
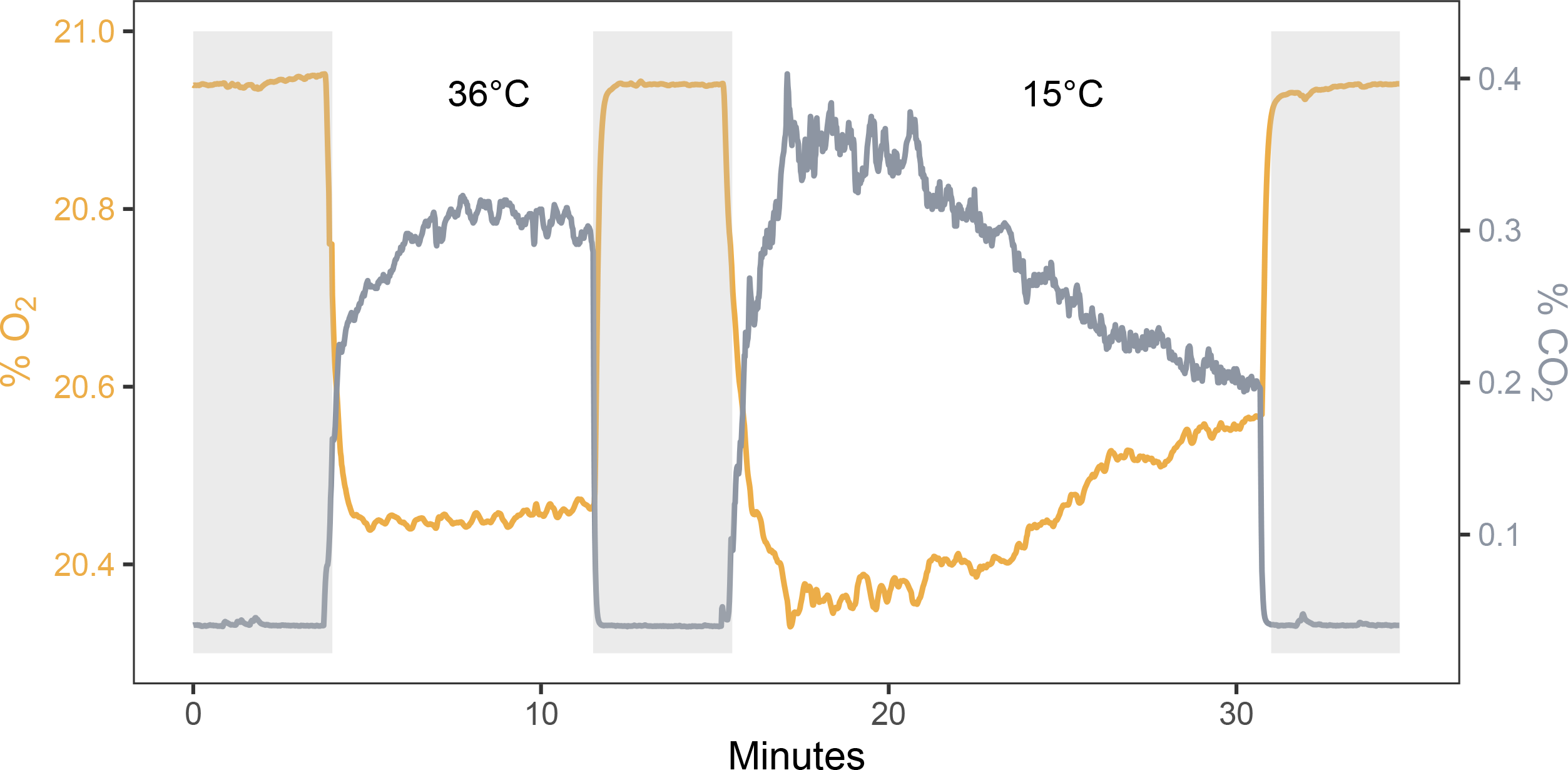
Representative respirometry data (nestbox 120 on May 28^th^ 2022) showing percent of O_2_ (orange) and CO_2_ (gray) in excurrent air during a full respirometry recording of 4 8-day-old nestlings first in a warm (36°C) and then a cold environment (15°C). Baseline measurements are shaded in gray.

We used equation 1 in (Lighton, 2018) to calculate oxygen consumption (VO_2_; mL min^-1^):

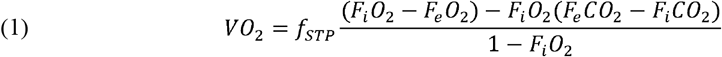

where F_i_O_2_, F_e_O_2_, F_i_CO_2_, and F_e_CO_2_ are the fractional concentrations (F) of incurrent (i) and excurrent (e) O_2_ and CO_2_. Data files were processed using ExpeData (v1.9.27). VO_2_ was converted to Watts (W; Joules per second) assuming an energy equivalence of 20 Joules per mL O_2_ (Kleiber, 1961). In the thermoneutral measurement, resting metabolic rate (RMR) was calculated as the mean of the most stable 2-min measurement of oxygen consumption. In the cold measurement, RMR was calculated as the rolling mean with the highest 30 s oxygen consumption. Based on these data, the metabolic response to cold temperature was calculated as:

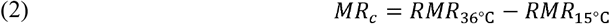

Cooling rates were calculated as the surface-area adjusted change in skin temperature during the first 5 min of cooling as in other studies (Andreasson et al., 2016; Page et al., 2022), according to equation 3,

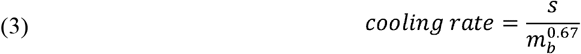

Where *s* is the slope of change in skin temperature (°C min^-1^), and *m*_*b*_ is body mass (in g). Body mass raised to the power of 0.67 was used to account for the scaling exponent between surface area and volume as in Andreasson et al. (2016) and Page et al. (2022).

### Data analyses

We used R for all data analyses (R Core Team, 2022). We used linear mixed-effect models with nest ID as a random intercept to evaluate our data using the R-package *lme4* (Bates et al., 2015). Assumptions of all models were assessed by visually inspecting residual plots and formal tests of multivariate normality using the *mvn* package (Korkmaz et al., 2014).

Body mass and cooling rate were both analyzed in models with age and brood size as categorical, independent variables. RMR_15°C_, RMR_36°C_ and MR_c_ were analyzed in similar models, with age and brood size category as factors and body mass (mean centered by age) as a covariate. We evaluated the full models including all possible interactions fitted with ML and type III sums of squares. Non-significant interactions were removed in a step-wise fashion based on F-tests using Satterthwaite’s method with a significance threshold of p ≤ 0.05 (Table S1). We then re-fitted final models using REML and used the R-package *emmeans* to calculate parameter estimates and their standard errors using the emmeans() function and predicted slopes with confidence intervals using the emtrends() function (Lenth, 2022). We controlled for repeated measurements in all models by including nest ID as a random effect.

Eleven nests were enlarged and 15 were reduced. However, during the course of the measurements, the sample size of reduced nests became smaller due to nest failure. One enlarged nest had to be excluded from the dataset on day 4 because the drying columns were not attached to the incurrent air valves. Skin temperature measurements were excluded when the thermocouple either fell off or broke during measurements. Resulting sample sizes are in Table 1.

**Table 1.** Number of broods of 4, 6, and 8-day old nestlings in experimentally enlarged (E) and reduced (R) nests included in estimates of metabolic rate and cooling rate.

| Age | Treatment | Metabolic rate: n | Cooling rate: n |
| --- | --- | --- | --- |
| 4 | E | 10 | 7 |
|  | R | 15 | 8 |
| 6 | E | 11 | 7 |
|  | R | 13 | 10 |
| 8 | E | 11 | 8 |
|  | R | 12 | 12 |

## Results

### Body mass

Body mass increased at similar rates in the two brood categories as they aged (age × treatment interaction: p = 0.83; main effect of age: *p* < 0.001) and body mass did not differ between brood size categories (p = 0.52; Table 2, Figure 2). When nestlings were 14 days old, we found no statistical difference between reduced and enlarged broods in body mass (t-test, t_20_ = -0.78, p = 0.44), wing length (t-test, t_20_ = 1.0, p = 0.33) or tarsus length (t-test, t_20_ = -0.56, p = 0.58).

**Table 2.**
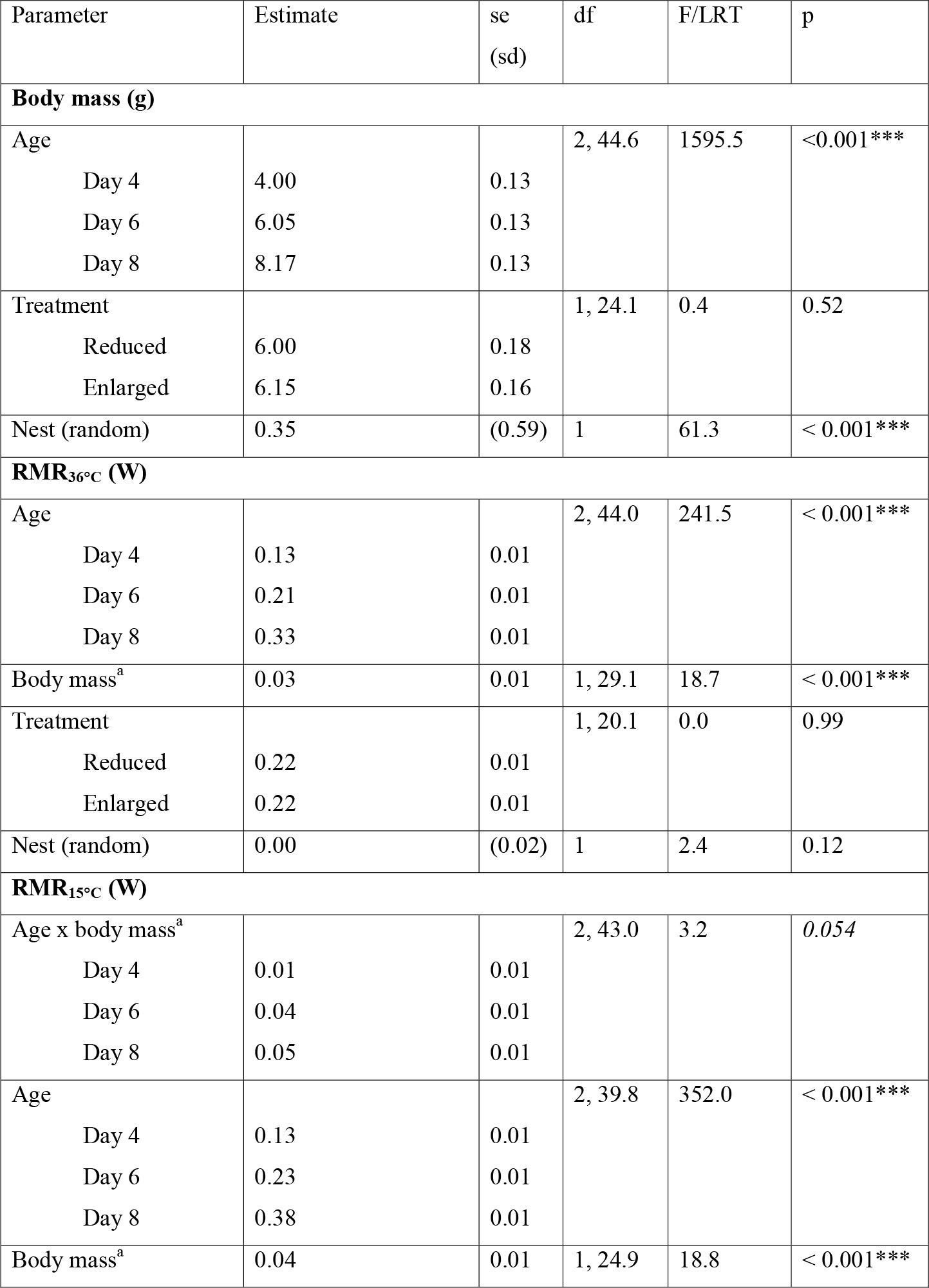
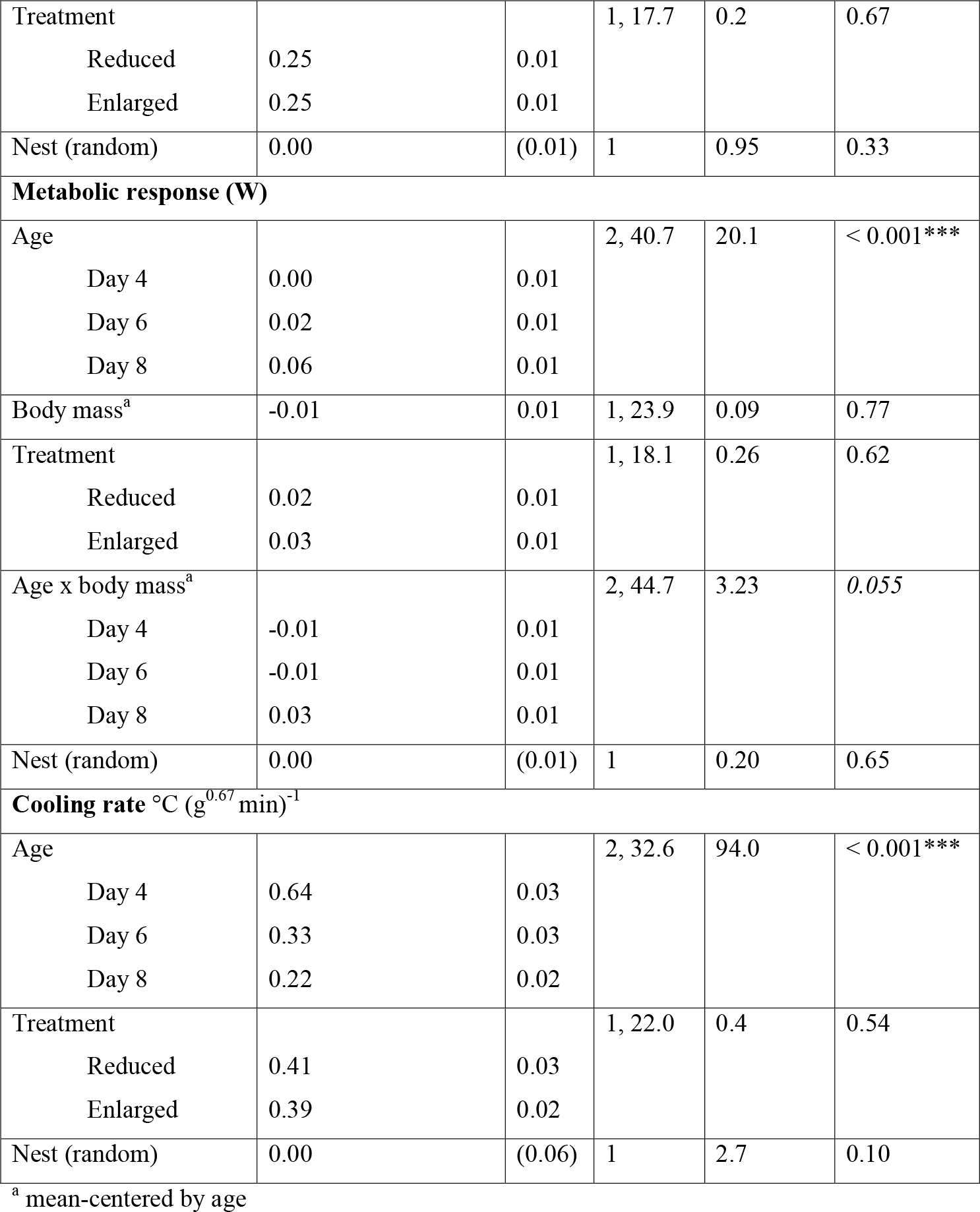
Results of Linear Mixed-effects Models (LMM) of nestlings’ body mass and metabolic response to a cooling challenge, and cooling rate on days 4, 6 and 8 after hatching and originating from experimentally enlarged or reduced broods. Parameter estimates (estimated marginal means or variance for random effects) are shown as mean values for age and treatment and as slopes for group mean-centered mass. Standard error, degrees of freedom, test statistics (fixed effects: F, random effects: LRT), and levels of significance are shown.

**Figure 2.**
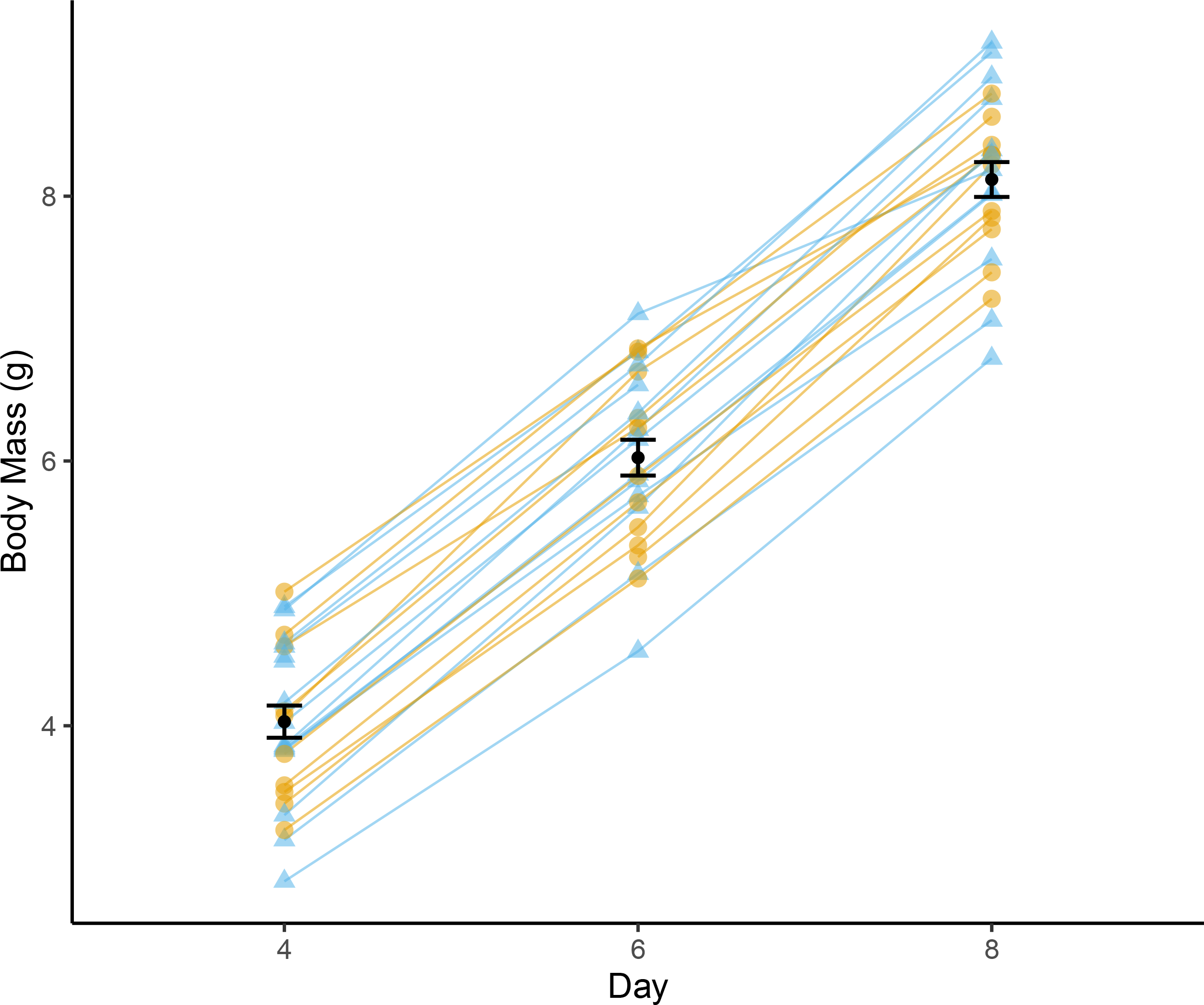
Body mass (g) of nestling blue tits on days 4, 6, and 8 and originating from enlarged (yellow triangles) or reduced (blue circles) broods size categories. Black points and bars represent mean and SE

### Resting Metabolic Rate

RMR_36°C_ increased with both age (p < 0.001) and body mass (p < 0.001, Figure 4), but did not differ between treatment groups (p = 0.99, Table 2). Averaged over body mass, there was a 62% increase between days 4 and 6 and a 57% increase between days 6 and 8 (estimated marginal mean +-SE: day 4: 0.13 +-0.01W, day 6: 0.21 +-0.01W, day 8: 0.38 +-0.01W, all pairwise comparisons using Tukey’s test were significant, p < .001). RMR_36°C_ increased by 0.033W per gram of body mass.

RMR_15°C_ increased with mass and there was a marginally significant increase in slope with age from 0.01 W/g on day 4 to 0.04 W/g on day 6 to 0.05 W/g on day 8 (age*mass interaction: p = 0.054, Table 2, Figure 4). There was also a positive overall effect of both mass and age on RMR_15°C_ (main effect of mass: p < 0.001; main effect of age: p < 0.001). There was a 77% increase in RMR_15°C_ between days 4 and 6 and a 65 % increase between days 6 and 8 (all pairwise comparisons using Tukey’s method were significant, p < 0.001; estimated marginal means and SE are shown in Table 2). We found no difference overall between enlarged and reduced broods in RMR_15°C_ (p = 0.66).

### Metabolic response to cooling challenge

The main effect of mass on metabolic response was marginally age-dependent (body mass*age interaction: p = 0.055, Figure 4) but we did not find a significant difference in slope in pairwise comparisons between ages (Tukey’s method; day 4 – 6: p = 0.89, day 4 – 8: p = 0.06, day 6 – 8: p = 0.1) and there was no significant relationship between mass and metabolic response when tested separately within age groups (linear regression; day 4: F_1, 23_ = 1.56, p = 0.22; day 6: F_1, 22_ = 0.23, p = 0.63; day 8: F_1, 21_ = 2.90, p = 0.10). The response was not significantly different from zero on day 4, but it was significantly different from zero on days 6 and 8 (paired t-test; day 4: t_24_ = -0.3, p = 0.77; day 6: t_23_ = 4.6, p < 0.001; day 8: t_22_ = 6.1, p < 0.001) and increased with age from 0.00 on day 4 to 0.02 on day 6 and 0.06 on day 8 (Table 2, Figure 3). We found no overall effect of either body mass (main effect of body mass: p = 0.77) or brood size category (main effect of treatment: p = 0.62)

**Figure 3.**
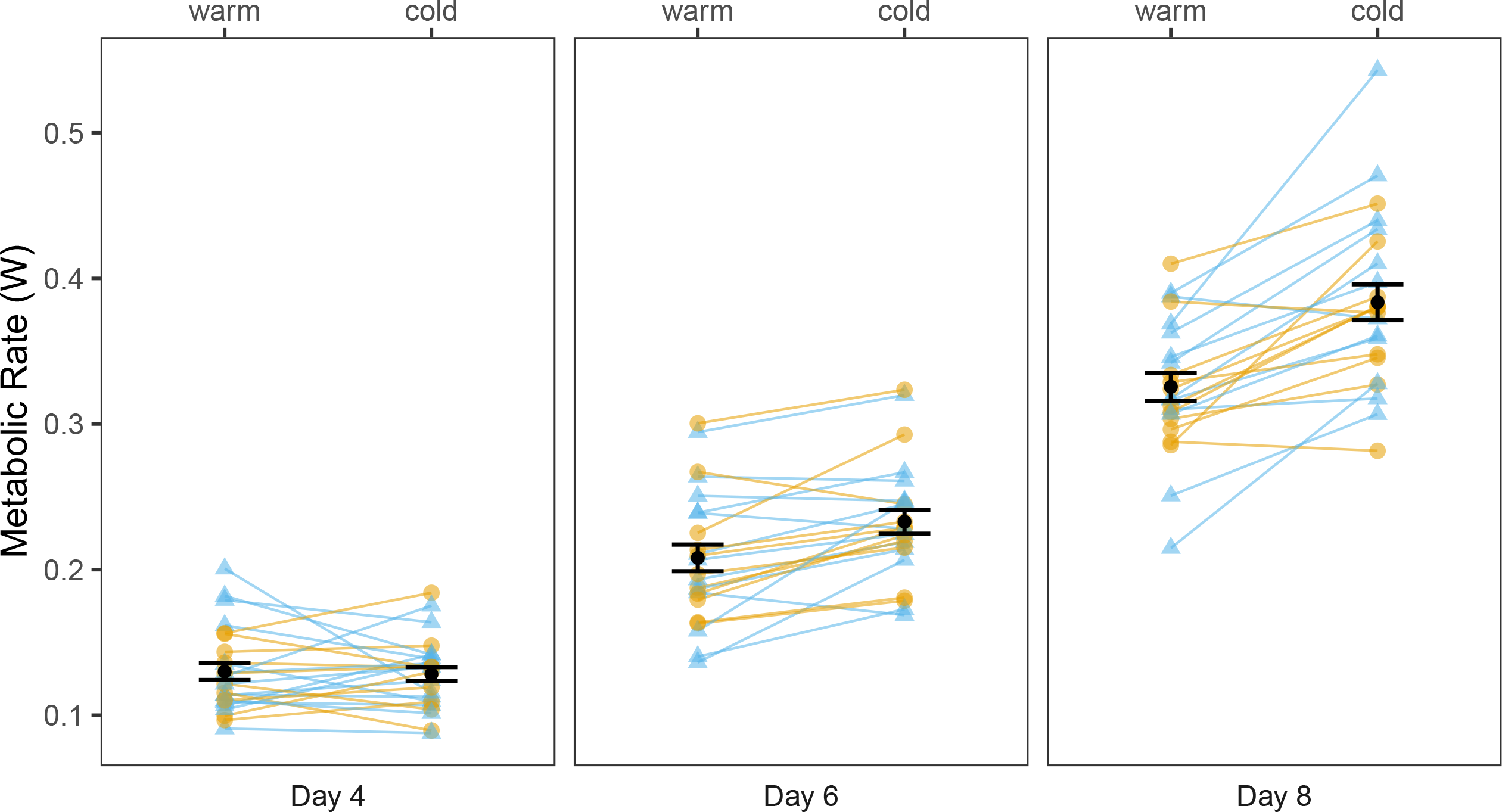
Nestlings from experimentally enlarged (yellow circles) and reduced (blue triangles) nests when 4, 6, and 8 days old showing the resting metabolic rate (W) of nestlings measured in a warm environment (36°C) and subsequent cold challenge (15°C). Black points and error bars show mean and SE.

**Figure 4.**
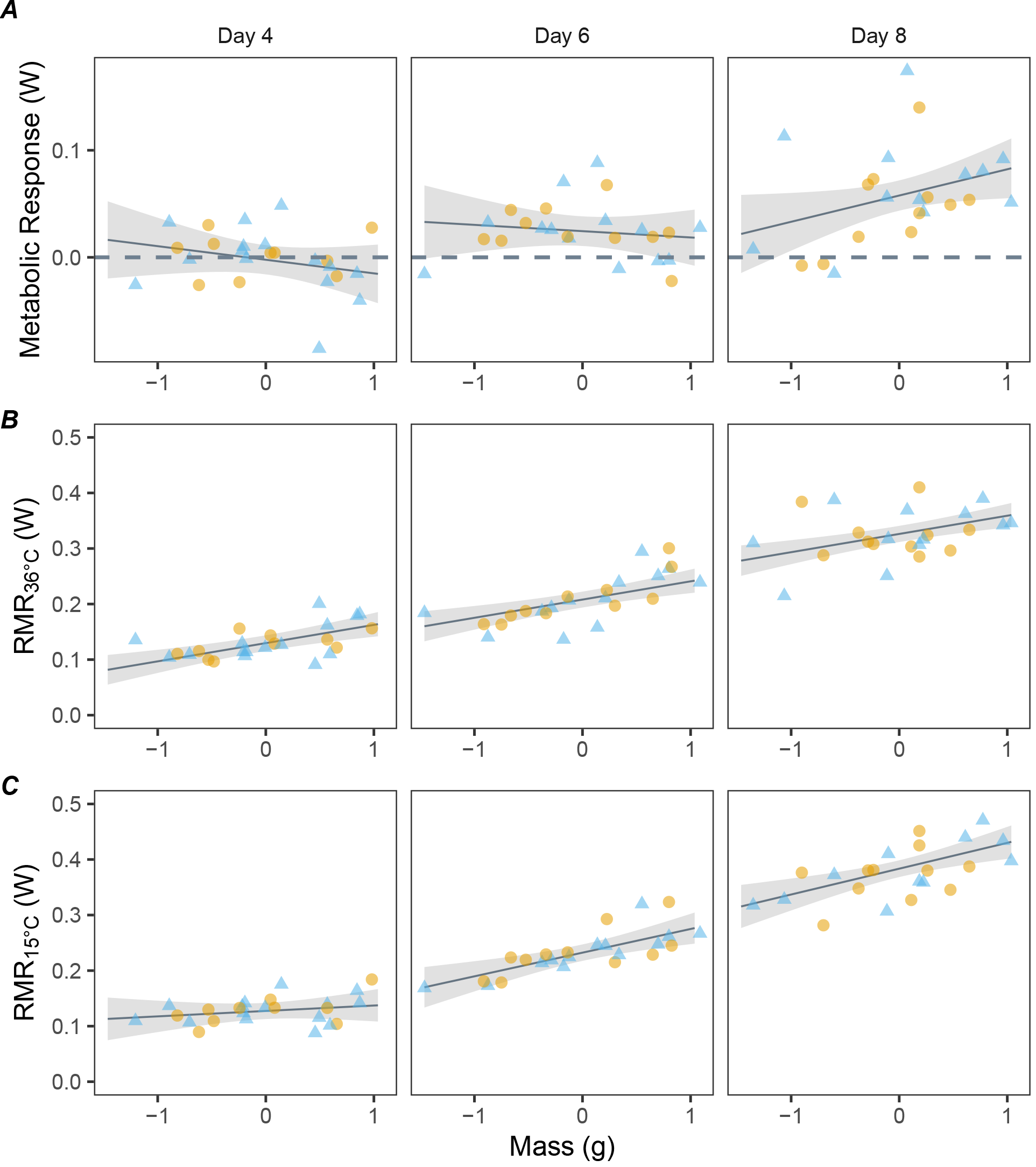
Metabolic traits of 4, 6, and 8-day-old nestlings in experimentally enlarged (yellow circles) or reduced (blue triangles) broods: A) metabolic response (maximum cold metabolic rate – RMR) with reference line at 0 (no response), B) RMR measured in 36°C, and C) RMR measured in 15°C in relation to group-mean-centered mass (±1.5 grams from mean mass (g)). Model predictions are shown as best fit lines with 95% confidence intervals.

### Cooling rate

Cooling rate decreased as nestlings grew older (p < .001) by half from day 4 to 6 and by a third from day 6 to day 8 (day 4: 0.64 +-0.03°C g^-0.67^ min^-1^, day 6: = 0.33 +-0.03°C g^-0.67^ min^-1^, day 8: 0.22 +-0.02°C g^-0.67^ min^-1^, mean +-SE; all pairwise comparisons using Tukey’s method significant, p < .001). We did not find a difference in cooling rate between the two brood size categories (p = .54, Table 2, Figure 5).

**Figure 5.**
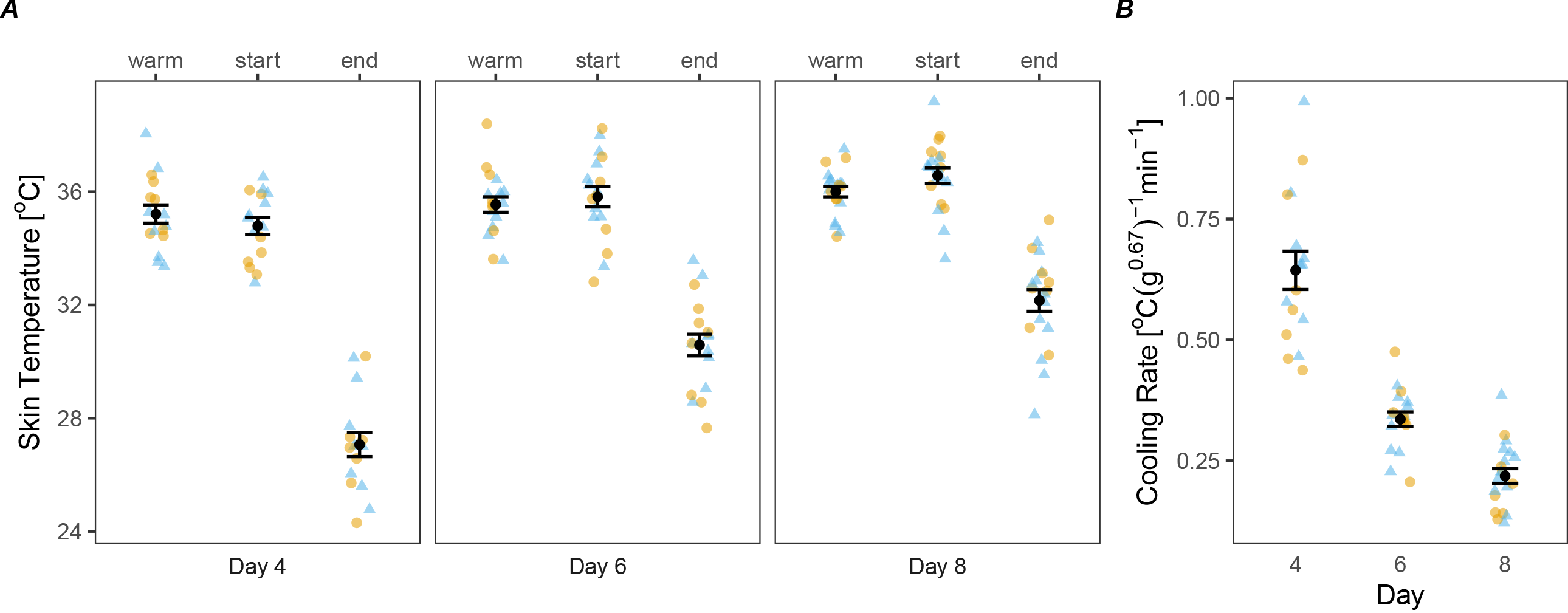
Nestlings from experimentally enlarged (yellow circles) or reduced (blue triangles) broods at age 4, 6, and 8 days with A) the mean skin temperature during a two-minute RMR measurement in 36°C (warm) followed by the temperature at the start and end the first 5 minutes of a 15°C cold-challenge (start, end). Panel B) shows the cooling rate during the first 5 minutes of the cold challenge. Black points and bars represent mean and SE.

## Discussion

We predicted that by experimentally manipulating brood size, we would change the amount of food received per nestling, with nestlings in larger broods receiving fewer food deliveries, as has been found previously in the same study system (in marsh and blue tits (Nilsson, 2002; Stjernman et al., 2004)). Based on this, we hypothesized that nestlings from enlarged broods would show a later onset of the development or use of endothermy compared to reduced broods, as was suggested by Andreasson et al. (2016). However, we found no differences in either RMR in 36°C (thermoneutrality) or RMR, metabolic response or cooling rate in a 15-minute, 15°C cooling challenge. Thus, we did not find support for our hypothesis during this year. Likewise, we found no differences in body mass, wing length, or tarsus length of nestlings in this study, despite a 36% change in mean brood size in both enlarged and reduced nests after brood size manipulations. One possible explanation is that we failed to manipulate the variation in food deliveries per nestling between the experimental brood categories. Previous studies show that while brood size enlargements usually result in lower quality nestlings in terms of reduced body mass, wing length, and/or tarsus length, the effects can be year-dependent (Nilsson & Gårdmark, 2001), particularly in response to years with a high resource abundance (Simon et al., 2004). A brood size manipulation study of great tits (*Parus major*) performed within 10 km of our study area in the same season (2022) also failed to find any effect on nestling size (personal communication, J.-Å. Nilsson). We believe that the lack of difference between the enlarged and reduced broods could be explained by such a year effect. Although we did not measure nest temperature in this study, our results also do not indicate any differences in the development of endothermy due to possible differences in the thermal environment of the nest attributable to brood size. In the remainder of the discussion, we focus on the overall development of endothermy in individual altricial nestlings.

### Metabolic response

We found that the first signs of endothermy in blue tits, a hole-nesting bird, coincide with the timing in other altricial birds, around days 4-6, (Breitenbach & Baskett, 1967; Clark, 1982; Dawson & Evans, 1957, 1960; Marsh, 1979; Mertens, 1977; Morton & Carey, 1971; O’Connor, 1975; Olson, 1992, 1994)). In our study, the mean metabolic response on day 4 was zero. The response became stronger as nestlings aged, with RMR_15°C_ being upregulated relative to RMR_35°C_ by 9% on day 6, and by 15% on day 8. At all ages, there was considerable individual variation in the metabolic response. This suggests that the onset of endothermy varied between nests due to factors unrelated to the experimental manipulations.

Both age and mass of nestlings are important predictors of metabolic rates. Thus, even rather small differences in mass within age groups would have an effect on heat production. When controlling for age, the effect of mass on the resting metabolic rates of nestlings was positive overall in both the cold and warm environment, but we did not find strong support for an age-dependent or overall effect of mass on the metabolic response. This is likely due to nestlings not yet having enough thermogenic tissues to mount a strong metabolic response, leading to their capacity being swamped quickly in the cooling experiment, regardless of their size. However, a positive trend seemed to emerge between body mass and metabolic response on day 8. It would be worth investigating if there is a gap in thermogenic capacity between nestlings of different mass that widens with age, especially considering that runts receive less food and grow at a slower rate compared to the rest of the brood.

The energy allocation hypothesis posits that altricial birds postpone the development of endothermy until after their most rapid growth period, to conserve energy for growth in the first days after hatching (Olson, 1992; Węgrzyn, 2013). Contrarily, blue tit nestlings, hole-nesting birds with a longer nestling period compared to their open-nesting counterparts, sustain a high growth rate up to day 10 (Andreasson et al., 2016, 2018), which is well after our observation of the onset of endothermy. However, none of the nestlings were able to sustain an elevated metabolic rate in 15°C by day 8 for more than a few minutes (Figure S2). This is likely due to nestlings having too little thermogenic tissue and having not yet developed their own insulation in the form of feathers. In other words, the heat produced by nestlings was not enough to account for the heat lost to the environment. Since the response in individual nestlings is brief up to at least day 8, any advantages of active thermoregulation would only have an effect during milder or shorter-term reductions in temperature in their immediate surrounding, such as when the female leaves the nest for short foraging bouts. Andreasson et al. (2016) measured nest temperature continuously and found that nest temperature increases and becomes relatively more stable after day 6 in outdoor nest boxes with a mean nightly ambient temperature of 13°C, while Mertens (1977) showed that great tits in the lab were able to maintain a stable body temperature in the nest in 15°C by day 6, with a brood size of 6. Therefore, for cooler and longer-term challenges, the ability to produce heat is probably only adaptive in the context of a brood, where nestlings are insulated by their siblings and nest materials.

### Cooling rate

In accord with the metabolic response, the cooling rate was best explained by the age of nestlings. We found that cooling rates were comparable to those in blue tit nestlings of the same age in other studies (Andreasson et al., 2016; Page et al., 2022). We observed from continuously recorded temperature data that skin temperature started to decline immediately after cold exposure and continued to decrease for the 15 minutes of measurement, so none of the nestlings were able to maintain a stable skin temperature in 15°C (Figure S2). However, cooling rates declined considerably with age (Table 2, Figure 5), and this is most likely due to a combination of an increase in resting metabolic rate, reduction in surface area to volume ratio, and increased capacity for a brief thermogenic response.

## Conclusions

The results of this study serve to improve our understanding of the development of endothermy in individual nestlings. However, to determine if and how brood size influences the ontogeny of endothermy, additional studies are needed to be carried out over multiple breeding seasons to control for potential year effects and could also benefit from including older nestlings and milder cooling challenges. Knowledge of the ontogeny of endothermy is important for our understanding of the thermogenic capabilities and the associated energetic requirements of nestlings at different stages of development, and to identify potential energy allocation tradeoffs between growth and development. This is especially true in a time when breeding phenology is advancing with the earlier arrival of spring, leaving birds more likely to experience opposing selection pressures such as cold snaps (Shipley et al., 2020) in the midst of higher mean temperatures during development (Andreasson et al., 2023).

## Acknowledgements

We would like to thank Johan Nilsson for his contribution to the fieldwork.

## Data Availability

Data will be available in the Dryad Digital Repository upon acceptance for publication

